# ADAM11 a novel regulator of Wnt and BMP4 signaling in neural crest and cancer

**DOI:** 10.1101/2023.06.13.544797

**Authors:** Ankit Pandey, Hélène Cousin, Brett Horr, Dominique Alfandari

**Affiliations:** University of Massachusetts Amherst, Department of Veterinary and Animal Sciences.

## Abstract

Cranial neural crest (CNC) cells are induced at the border of the neural plate by a combination of FGF, Wnt, and BMP4 signaling. CNC then migrate ventrally and invade ventral structures where they contribute to craniofacial development. Here we show that a non-proteolytic ADAM, Adam11, originally identified as a putative tumor suppressor binds to proteins of the Wnt and BMP4 signaling pathway. Mechanistic studies concerning these non-proteolytic ADAM lack almost entirely. We show that Adam11 positively regulates BMP4 signaling while negatively regulating β-catenin activity. By modulating these pathways, Adam11 controls the timing of neural tube closure and the proliferation and migration of CNC. Using both human tumor data and mouse B16 melanoma cells, we further show that ADAM11 levels similarly correlate with Wnt or BMP4 activation levels. We propose that ADAM11 preserve naïve cells by maintaining low Sox3 and Snail/Slug levels through stimulation of BMP4 and repression of Wnt signaling, while loss of ADAM11 results in increased Wnt signaling, increased proliferation and early epithelium to mesenchyme transition.

## Introduction

The development of neural crest cells during embryogenesis involves both pathways that regulate cellular migration and invasion which are comparable to the most aggressive tumors (Medina-Cuadra and Monsoro-Burq, 2021). Often, tumorigenesis involves a first step in which cells lose their fully differentiated state to revert to a more naïve pluripotent state. In addition, once the original tumor is formed, the cells may undergo epithelium to mesenchyme transition to evade the original tissue and invade a new target site where they create metastasis (Chaffer et al., 2016; Wilson et al., 2020). Understanding the molecular underpinnings of this transition is critical for developing strategies to stop it in tumors (Shibue and Weinberg, 2017).

Here, we investigated the role of non-proteolytic ADAM proteins in EMT and cellular migration in the cranial neural crest cells. In the embryo, the cranial neural crest cells are induced at the border of the neural and non-neural ectoderm by signals emanating from the surrounding tissues that include Wnt, FGF, and BMP4 (Bastidas et al., 2004; de Croze et al., 2011; Garnett et al., 2012; Steventon et al., 2009). Once induced, the CNC cells remain stem-like and progress through an EMT to migrate and invade the ventral structures of the embryo where they differentiate into bone cartilage and ganglia of the face (Alfandari et al., 2010; Scarpa et al., 2015).

While other ADAMs have a well-established role in CNC migration (Alfandari et al., 2001; Cousin et al., 2012; Khedgikar et al., 2017; McCusker et al., 2009; Neuner et al., 2009), the role of non-proteolytic ADAMs, which are also expressed in this tissue, during development has not been studied. These ADAMs are single-pass transmembrane proteins that possess the same domain organization as other members of the family, including a pro-domain, metalloprotease domain, disintegrin and cysteine-rich domain, EGF domain, and a short transmembrane and cytoplasmic domain. What makes them stand out is that the key glutamic acid in the protease active site is not present rendering them proteolytically dead (Hsia et al., 2019). Proteolytic ADAMs control Notch, EGF, TGF, Ephrin, and Wnt signaling pathways by cleaving substrate or receptors to activate or inhibit the relevant signaling pathways (Alfandari et al., 2009; Klein and Bischoff, 2011; Reiss and Saftig, 2009). In contrast, non-proteolytic ADAMs have been shown to interact with multiple transmembrane proteins including integrins, LGI, and Kv channels, and to play significant role in synapse organization that establishes sensory neural circuits (Hsia et al., 2019; Sagane et al., 2008; Wang et al., 2018).

We have previously shown that *Adam11* is expressed in the cranial neural crest and a subset of neurons in the early neural tube in *Xenopus laevis* (Cai et al., 1998). A very similar expression pattern was described in the mouse embryo (Diez-Roux et al., 2011; Rybnikova et al., 2002). Mice lacking *Adam11* are viable yet have deficiencies in pain perception suggesting a key role for the protein in establishing the normal neuronal connections responsible (Takahashi et al., 2006a; Takahashi et al., 2006b). Here we show that loss of Adam11 during early embryogenesis results in early neural tube closure and early EMT of the CNC *in vitro*. We further show that Adam11 promotes BMP4 signaling and reduces Wnt/β-catenin signaling *in vivo*. The Wnt signaling pathway has been directly involved in the increased proliferation through control of *CyclinD1* (Kafri et al., 2016; Koehler et al., 2013) and EMT with the increase in *Snail* transcription factors expression driving the loss of epithelial cadherins and the progression of EMT (Yook et al., 2005). Loss of Adam11 in embryos results in an increased β-catenin activation leading to premature EMT, higher expression of *Snai2* and *CyclinD1*, and increase CNC proliferation.

Interestingly, decrease in *ADAM11* expression is found in the majority of solid tumor that have been studied and correlates with an increase in Wnt signaling and *CYCLIND1* expression (this study). Decreasing Adam11 in B16 mouse melanoma cells similarly increases cell proliferation by promoting the G1-S transition, a step controlled by CyclinD1. We also show that Adam11 increases BMP4 signaling resulting in increased Smad1-5 phosphorylation and a decrease in Sox3 expression suggesting that Adam11 reduces Sox3 expression in the naïve CNC to prevent their early differentiation. Similarly, in the neural crest derived tumor, neuroblastoma, *ADAM11* expression is higher than in the normal tissue, with a concurrent increase in *Smad1-5* expression and decreases in *Sox3* expression.

We propose that Adam11 maintains stem cell like attributes in both CNC and cancer and that either increase in Adam11 or a decrease in Adam11 can create an unbalance toward stemness or EMT contributing to the cancer pathogenicity.

## Results

### Adam11 Knockdown (KD) accelerates neural tube closure.

*Adam11L* is expressed in the central nervous system and the cranial neural crest (Cai et al., 1998). The mRNA for both S and L allo-alleles is expressed zygotically at low levels. *Adam11S* is the major form expressed during gastrulation, while *Adam11L* is the main form during neural tube closure and neural crest cell migration (Xenbase RRID:SCR_003280, st17 to 25) (Fortriede et al., 2020; Session et al., 2016). To test the role of Adam11 we designed two sets of morpholino to either block translation of both the *Adam11L* and *S* allo-alleles (MO11, L and S) or block the splicing of the first intron of *Adam11.L* which is the main form expressed within the cranial neural crest cells (MO11spl). The efficiency of these morpholinos was tested by western blot (Fig.1A) and RT-qPCR (Fig.1C). We used time-lapse imaging to follow potential developmental defect after the onset of zygotic transcription (Stage 8). While we did not see defect during gastrulation, we observed that the neural tube closed earlier in morphant embryos when compared to control (Fig.1C). This was true for both splicing and translation blocking morpholino and is a very rare phenotype as injections typically result in a slight delay of development but rarely an apparent acceleration (DA personal observation). We were able to rescue this phenotype by injecting Adam11L mRNA lacking the 5’ untranslated region (Adam11L_R), a form that is not affected by the morpholino (Fig.1C), confirming that the phenotype was indeed specific (Fig.1C). Given that both Morpholinos gave the same phenotypes we focused for the rest of the study on the translation blocking morpholino that blocked both allo-alleles.

**Figure 1:**
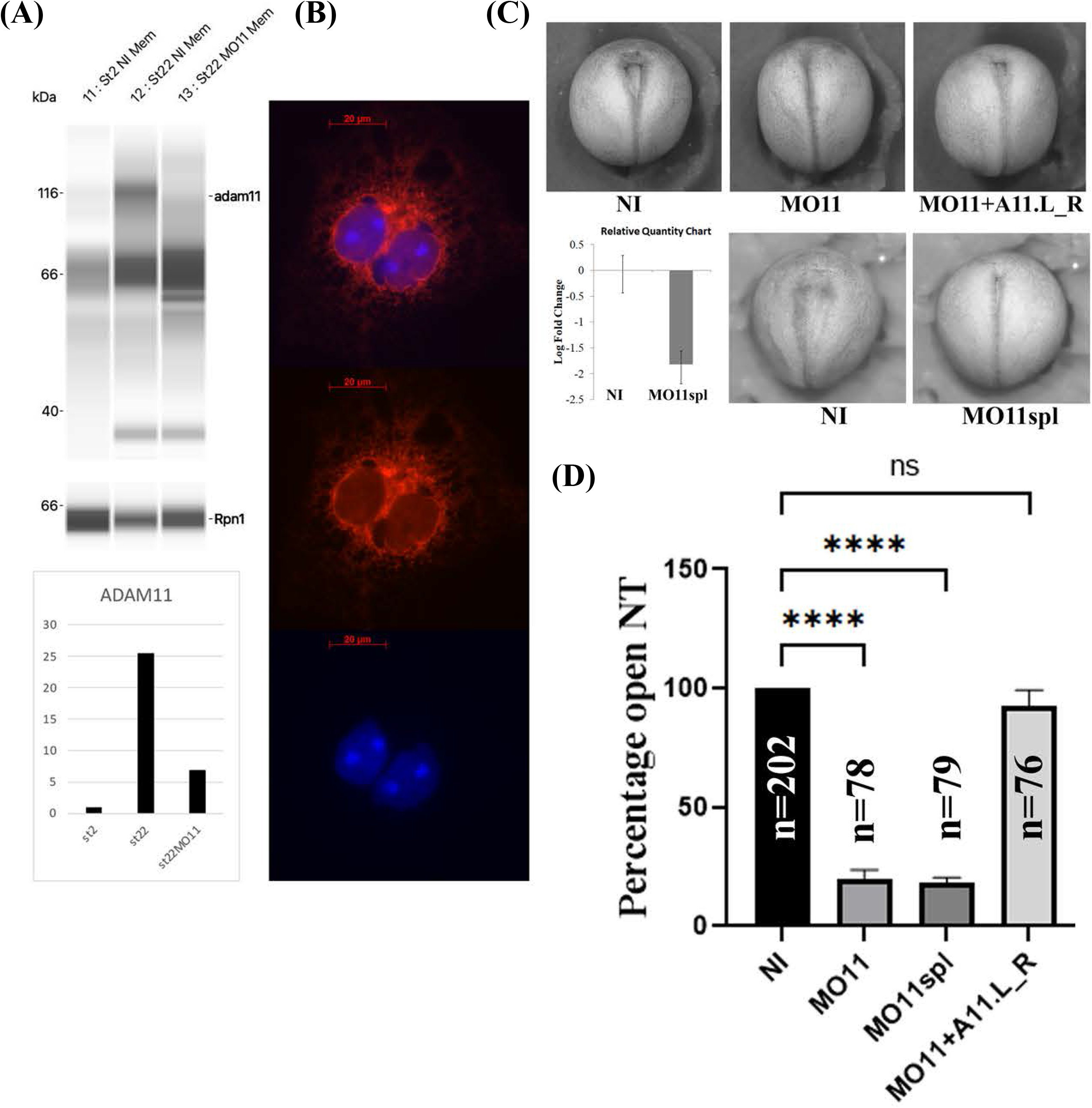
Adam11 Knock Down. A) Western blot with a monoclonal antibody (mAb) to Adam11 (DA3C5) using membrane extract from 10 embryos at stage 2 and 22. Knockdown efficiency of a translation blocking MO for Adam11 (MO11) was tested on either non-injected control embryos or embryos injected at the one-cell stage with 10 ng of MO11. The histogram depicts the quantification of Adam11 protein amount normalized to ribophorin 1 (rpn1) in each lane. (B) Immunofluorescence image of Adam11 staining using the mAb 3C5 on cos7 cells transfected with Adam11L. (C) Representative photograph of neurula stage embryos (Dorsal view). Anterior is up. Embryos were injected at the one-cell stage with either the translational blocking MO11 or Splicing blocking MO11spl. Embryos injected with MO11 were rescued using 12.5pg of ADAM11_R rescue mRNA. lacking the morpholino binding sequence. The histogram represents RT-qPCR from embryos injected with Adam11.L oligonucleotides designed to only amplify the spliced mRNA. D) Histogram representing statistical analysis of the phenotype represented in D. Asterisk represent the statistical significance (ANOVA) at p<0.0001 (****).

### Loss of Adam11 affects neural crest cell migration

Because of *Adam11L* expression in the cranial neural crest cells (CNC), we tested the ability of CNC with reduced Adam11 protein to migrate *in vivo* and *in vitro* (Fig.2). *In vivo* we tested the migration capacity by targeted injection (Fig.2A-B) as well as grafting experiments (Fig.2D-E) using previously described assays (Abbruzzese et al., 2016; Cousin et al., 2011; Cousin et al., 2012). In both experiments, we found that the loss of Adam11 perturbed CNC migration significantly in approximately 50% of the embryos (Fig.2B and 2E). In the grafting experiments we found significant defect in the CNC migration into the Branchial segments but not in the mandibular pathway. The inhibition of CNC migration was rescued by injection of the *Adam11L* rescue mRNA in the CNC precursor cells (Fig.2B). As we have shown before, inhibition of CNC migration *in vivo* can be due to either a loss of cell motility (Alfandari et al., 2003), or a more complex effect of the cellular environment *in vivo* (Cousin et al., 2012). To test this possibility, we placed CNC explants *in vitro* on fibronectin coated substrates and performed time-lapse video microscopy. Using this assay, we have previously shown that CNC explant dissected at stage 17 migrates collectively during the first 5 hours before individual cells break away, a process also observed *in vivo* (Alfandari et al., 2003; Alfandari et al., 2001). We found that CNC cells in both control and Adam11 KD were able to spread and migrate on the substrate efficiently, but that cells lacking Adam11 migrated individually even during the first 5 hours (Fig.2F). Furthermore, we found that the average velocity and persistence of migration was higher in cells with reduced Adam11 (Fig.2G). Collectively these results show that Adam11 is essential for proper CNC migration and may regulate the timing of neural tube closure and single neural crest cell migration. We then tested if neural and neural crest cell marker were expressed and localized properly in embryos lacking Adam11.

**Figure 2:**
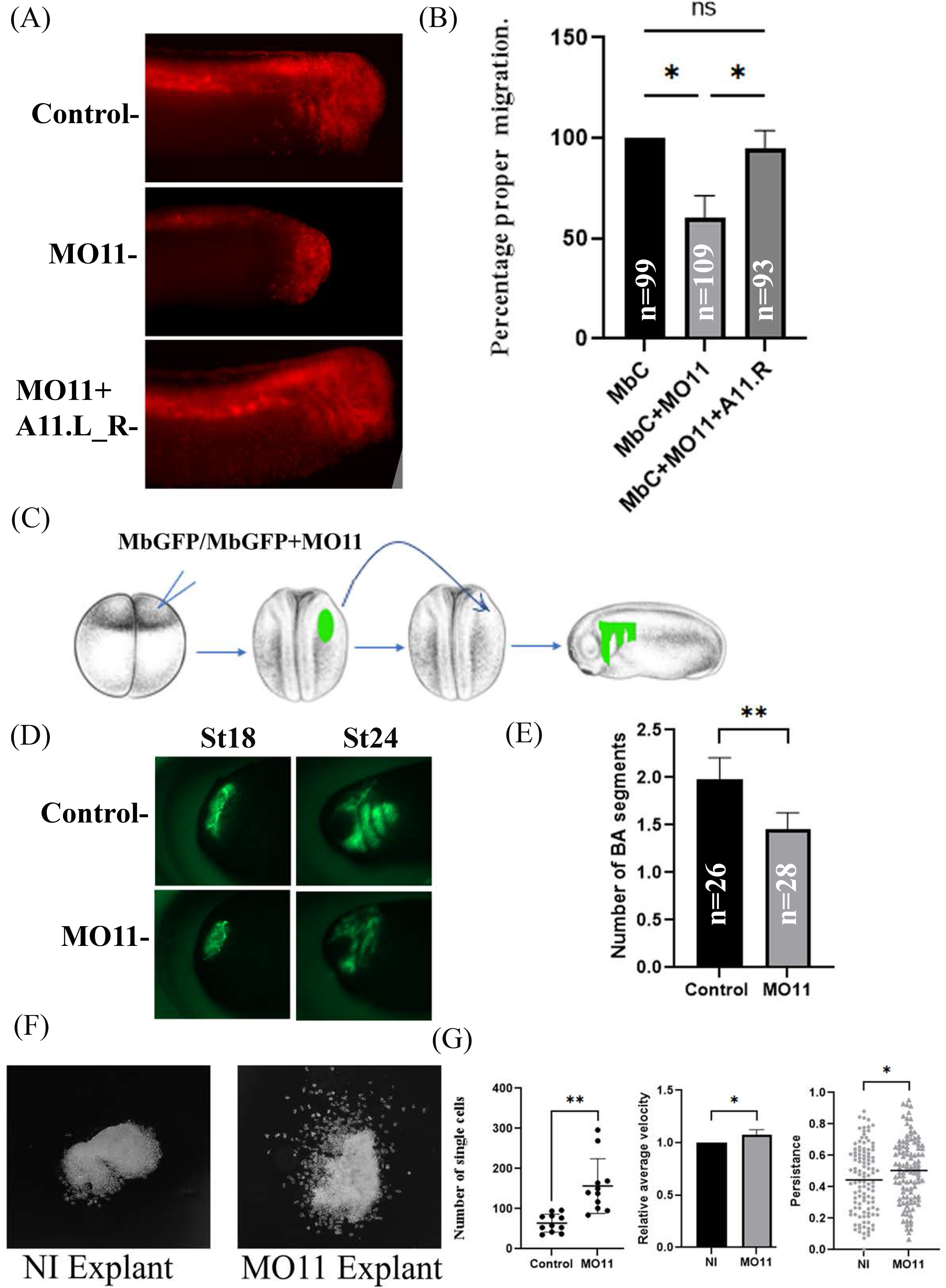
Adam11 Knock down affects cranial neural crest cell migration. A) Fluorescent tracking of CNC migration. Lineage tracer (Mb Cherry) was injected at the 8-cell-stage in one dorsal animal blastomere, either alone (control) or with the morpholino to Adam11 or both MO11 and the Adam11L_R rescue mRNA (125 pg). The presence or absence of fluorescent neural crest on the lateral and ventral side of the embryo is scored only in embryo with proper targeting (Dorsal anterior quadrant). B) Statistical analysis of the targeted injection. The percentage of embryos with fluorescent CNC in the migration pathway is represented (Y axis) normalized to Mb Cherry alone (100%). A minimum of 3 independent experiment was performed. Asterisks represent a p-value for ANOVA< 0.05. C) Schematic diagram of CNC grafting experiment. D). Representative photographs of embryo grafted with fluorescent CNC explant at stage 18 and scored at stage 24 for both control and MO11 injected embryos. E) Statistical analysis of the grafting experiment (n is the number of embryos). The histogram represents the number of branchial arches (BA) segments scored at st24stage 24 (**=P<0.01). F) Representative example of Cranial neural crest cell explants on fibronectin. Photographs were taken during the initial phase of collective cell migration (5 hr). G) Statistical analysis of cellular migration showing the number of single cells per explant, the relative average velocity and the persistence of cell migration. Single cells were scored at 5h. The average velocity was measured for 110 single cells over (30 minutes). The persistence of migration was measured on the same cells by dividing the distance from the start and end position by the distance travelled. (**= P<0.01, *=P<0.05).

### Loss of Adam11 affects neural and neural crest cell markers at the neurula stage

Given the distribution of Adam11L mRNA and the phenotypes observed following Adam11 KD, we focused our analysis on neural and neural crest markers at stages when the first phenotype is visible (Neurula stage). We used the well-characterized neural crest marker *Slug/Snai2* (Fig.3A), *Sox8* (Fig.3B) and *Sox9* (Fig.3C) in embryos injected unilaterally at the 2-cell stage with MO11. This allows KD in one half of the embryo (right or left side). *In situ* hybridization revealed that all three markers were expressed at the appropriate position. Interestingly, we found that these markers were increased on the injected side (Right and Fig.3F). Expansion of the neural crest territory can, in some cases, be done at the expense of the neural ectoderm. We tested the position of two neural markers, *Sox2* (Fig.3D) by *in situ* hybridization and Sox3 by immunostaining (Fig.3E). In both cases we found that the domain of expression on the injected side was also wider, suggesting that the increase in neural crest territory was not done as the expense of the neural tissue.

**Figure 3:**
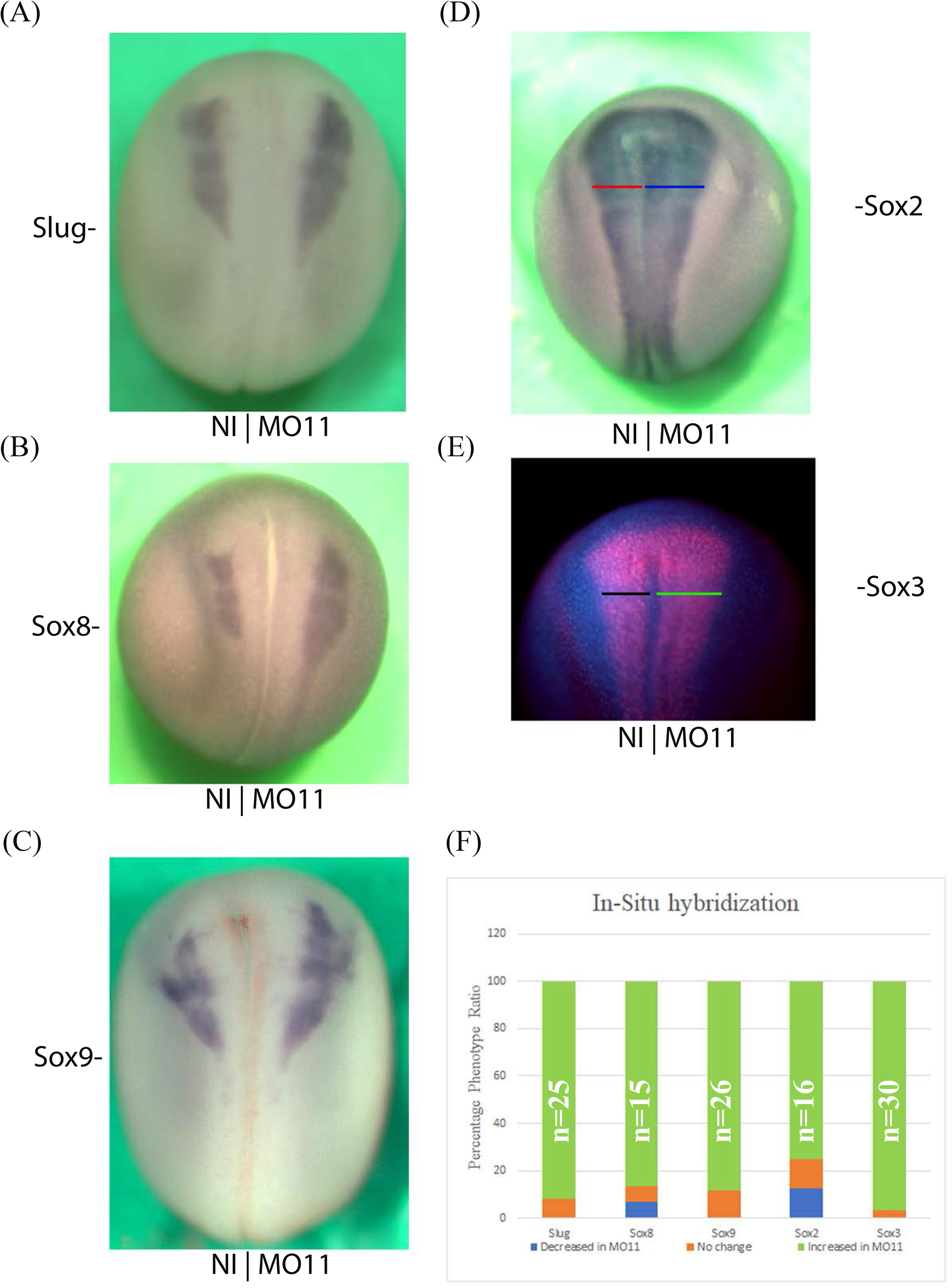
Adam11 KD affects neural and neural crest cell markers at neurula stage. Dorsal views of representative *in situ* Hybridization with neural crest (A-C) and Neural markers (D). The injected side is to the right, anterior is toward the top. (E) Immunodetection of the neural marker Sox3 with the same orientation as above. Colored lines in D and E highlight the increased size of the neural plate on the injected side. (F) Histogram depicting phenotype of the above-mentioned markers.

### Loss of Adam11 increases β-catenin activity in the CNC

Given the early onset of single CNC cell migration and the increase in *Slug* expression, we hypothesized that β-catenin controlled EMT could be dysregulated following Adam11 KD. To test this hypothesis, we dissected CNC explants from embryos injected at the two-cell stage in one cell and compared the explants from the control and the injected side after migration on fibronectin coated slides (Fig.4). Membrane Cherry was co-injected with the morpholino to identify the injected side and confirm that the CNC had been targeted. We found that while the overall fluorescence of β-catenin was similar on the control and injected side, the nuclear localization of β-catenin was expanded on the injected side (Fig.4A and 4C compare NI to MO11). In control explants, the β-catenin protein was present near the plasma membrane and in the nuclei of the cells that are at the leading edge of the explant, but was restricted to the plasma membrane in cells within the explant (Fig.4C, NI)(Alfandari et al., 2010). In contrast, explants missing Adam11, show nuclear β-catenin localization all the way through the explant (Fig.4C MO11). In addition, it is also clear that the nuclei of the control explants are more packed (Fig.4A DAPI) than the one from the Adam11 KD suggesting that the cells are more spread on the FN substrate. Nuclear β-catenin is known to activate gene expression including *slug* (Fig.3A) and *cyclin D1*. We, therefore, tested the expression of CyclinD1 in CNC explants (Fig.4B). Again, while the expression of CyclinD1 is mostly restricted to the periphery of the CNC explant from the control side, the expression is present throughout the explant from the injected side, and the overall intensity is significantly higher than in the control side (Fig.4D). It is important to note that these images are maximum projections so that protein expression levels should not be affected by their relative Z-position within the explant.

**Figure 4:**
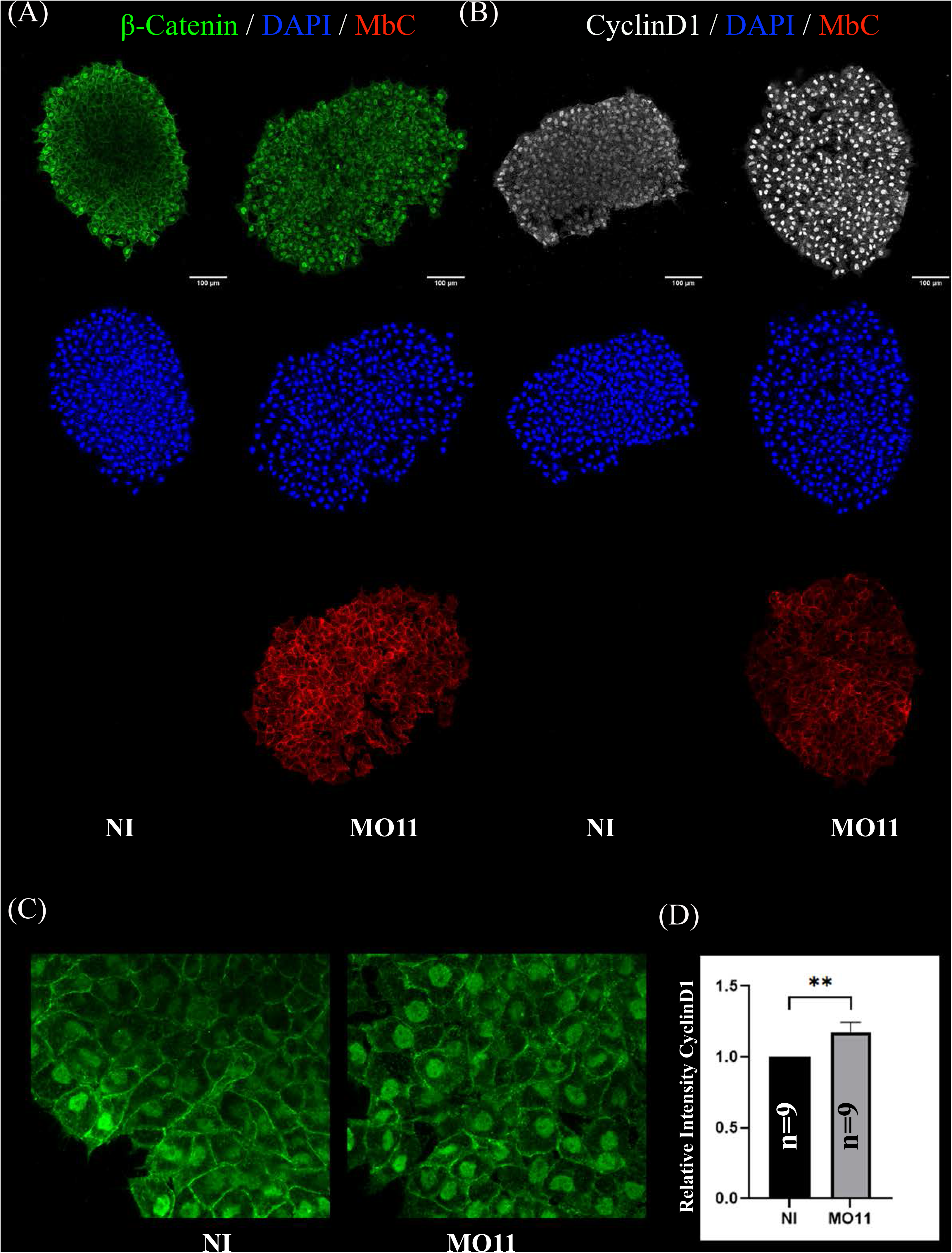
Adam11 KD increases β-catenin activity in the CNC. (A-C) Confocal imaging of CNC explants one hour after dissection. CNC explants were dissected at stage 17 and placed on fibronectin coated glass coverslip. Explants were stained for β-catenin (A and C, green), Cyclin-D1 (B, white) and nuclei (DAPI, blue). (C) Higher magnification of the β-catenin staining. Scale bars represent 100 µm. (D) Histogram depicting the relative intensity of CyclinD1 immunofluorescence between control CNC explants and Adam11 knockdown explants. Students t-test was done for statistical analysis (*=P<0.05).

### Loss of Adam11 leads to higher proliferation and early EMT

To directly test if the β-catenin transcriptional activity was affected by the loss of Adam11 we used the well-characterized Top-Flash luciferase reporter (Baarsma et al., 2011; Chuang et al., 2010) *in vivo*. To do this, we targeted the Top flash reporter to the precursor CNC territory at the 16-cell stage (Moody, 1987) in either control embryos or embryo KD for Adam11. We found a statistically significant increase in Top flash activity in embryos KD for Adam11 (Fig.5B).

**Figure5:**
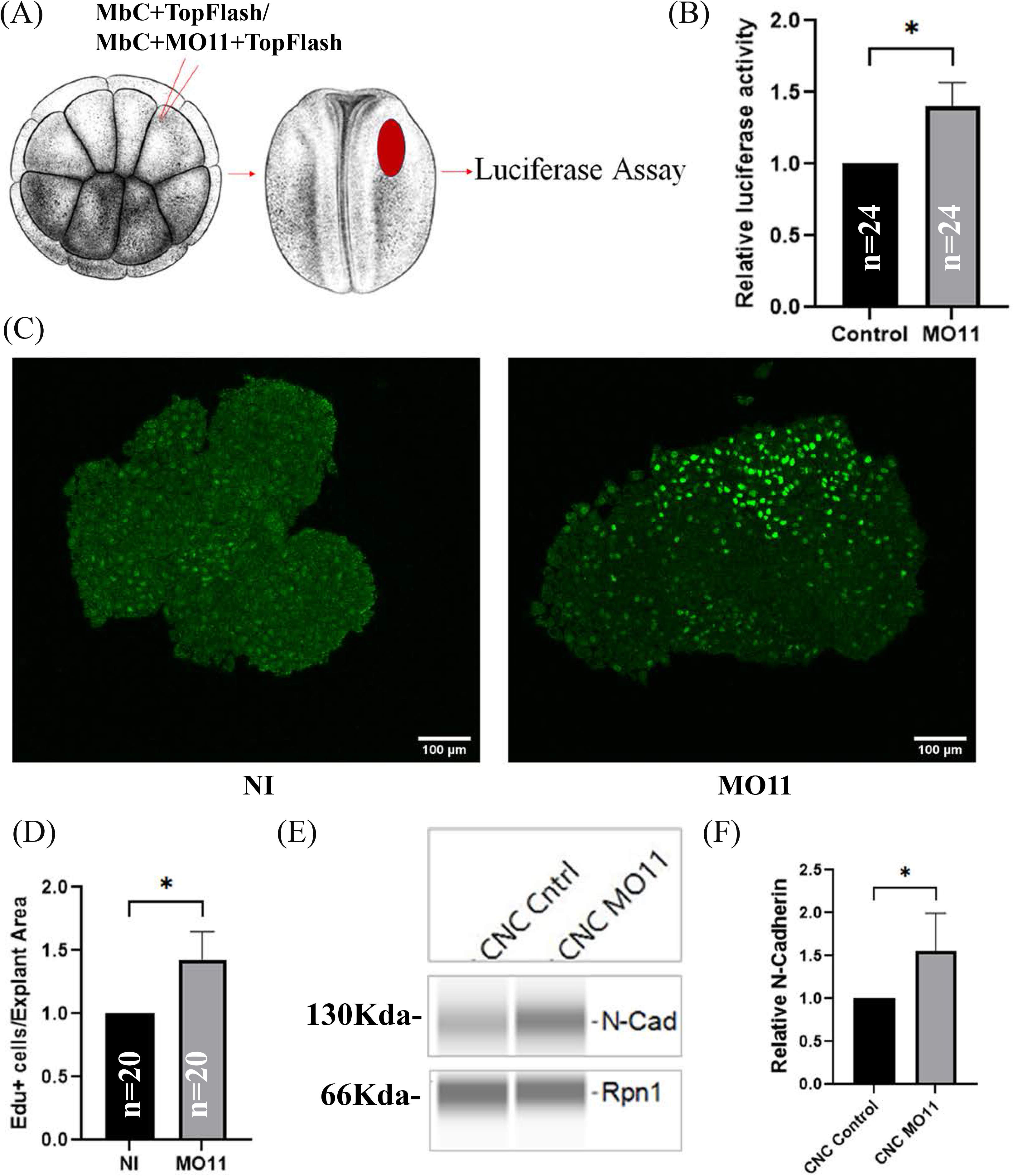
Adam11 KD leads to increase β-catenin transcriptional activity, higher proliferation and increased N-cadherin expression. A) Graphical representation of the luciferase experiment in embryos. B) Top flash luciferase reporter activity. 16-cell stage embryos were injected with a lineage tracer, and 10 pg of the top flash reporter plasmid together with 2 pg of the CMV renilla plasmid in one dorsal animal blastomere. Embryos with lineage tracer present in the dorsal anterior quadrant at neurula stage were selected and extracted individually for dual luciferase reporter assay. Control embryos were compared to embryos injected with MO11 (*= p<0.05). C-D) Quantification of CNC proliferation. C) Confocal imaging of Edu (green) labeled CNC explants three hour after dissection. Embryos were incubated with Edu at st15stage 15 for 2 hours. D) Histogram showing the relative number of Edu positive cells. Numbers were normalized to the size of each explant. (E) Capillary western Blot of N-cadherin from CNC explants dissected at stage 17. F) Histogram showing N-cadherin protein level normalized to Rpn1. Students t-test was performed for data represented in B, D, F (*=P<0.05).

Given the known role of CyclinD1 in controlling the G1 to S transition (Stacey, 2003), we tested the proliferation of CNC cells using Edu labeling. Embryos were incubated in Edu for two hours at the beginning of neurulation (stage 15/16), when the expression of Adam11 mRNA is maximum (Session et al., 2016). CNC explants were dissected at stage 17 and allowed to migrate on fibronectin for three hours prior to fixing and staining for Edu positive nuclei (Fig.5C). We observed a significant increase in Edu-positive cells in CNC with lower Adam11 when compared to control explants (Fig.5D). To account for variability in explant size and labelling stage between experiments, the number of EDU positive was normalized to the size of each explant and set to one for the control explants. Loss of E-cadherin and gain of N-cadherin is a classical hallmark of EMT in CNC and in cancer (Araki et al., 2011; Cousin, 2017; Huang et al., 2016; Scarpa et al., 2015), we therefore tested the expression of N-cadherin via quantitative capillary western blot on protein extracts from stage 17 CNC explants. When compared to Ribophorin 1 (Rpn1 as a loading control), we found that CNC lacking Adam11 expressed more N-cadherin than the control explants (Fig.5E-F). Taken together these results show that loss of Adam11 increases β-catenin transcriptional activity and causes earlier EMT, suggesting that Adam11 role is to reduce and/or delay this activity in the neural crest.

### Adam11 binds to protein from the BMP4 and Wnt signaling pathways

While the apparent early EMT pointed to β-catenin, we wanted to use an unbiased approach to identify additional functions for Adam11 in neural and neural crest cells. Adam11 is a non-proteolytic ADAM with no known role during embryogenesis. Protein interaction studies have identified putative binding partners for Adam11 in mouse including integrins and LGl proteins in neurons (Kole et al., 2015; Sagane et al., 2008; Takahashi et al., 2006a; Takahashi et al., 2006b; Wang et al., 2018). These putative protein interactors cannot explain the effect of Adam11 KD on β-catenin activation. In the absence of an antibody to Xenopus Adam11 we expressed the Flag-tagged Adam11 in Xenopus Cranial neural crest cells and human Hek293T cells and pulled down interacting proteins (Fig.6). In embryos, we injected at the 8 cell-stage to target the CNC and extracted the proteins at either the neurula or tailbud stages (4 samples from two independent experiments). We considered proteins with at least 2 unique high confidence peptides (95%) that were present in at least two of the 4 samples as potential candidates. We also performed Flag pull down on human Hek293T cells transfected with Adam11-Flag or RFP-Flag. Among the proteins that were found in both data sets, we identified the secreted protein BMP4 as well as the β-catenin transcriptional co-activator Bcl9. We also found multiple members of the LRP family of protein, and the Wnt receptor FzD6 but these were not present in both data sets (LRP2 in embryos, LRP5-6, and Fzd6 in Hek293T cells). We also found the BMP4 receptor BMPR1A in Hek293T cells. Given the critical role of the Wnt/β-catenin and BMP4 signaling pathway to both neural and neural crest cell development and our results showing increased β-catenin activity, we focused on these putative binding partners for this study. In addition, other ADAM have been shown to interact with Wnt receptors or small secreted forms of the receptor that act as antagonists (Abbruzzese et al., 2015; Esteve et al., 2011), thus we also tested the interaction with Wnt receptor Fzd. We were able to confirm the interaction of Adam11 with BMP4 (Fig.6C) and the BMP receptor 1A (Fig.6D) by co-immunoprecipitation. We also identified the interaction of Adam11 with the Wnt receptor Fzd7 a key component of the β-catenin signaling pathway in embryos (Fig.6E).

**Figure 6:**
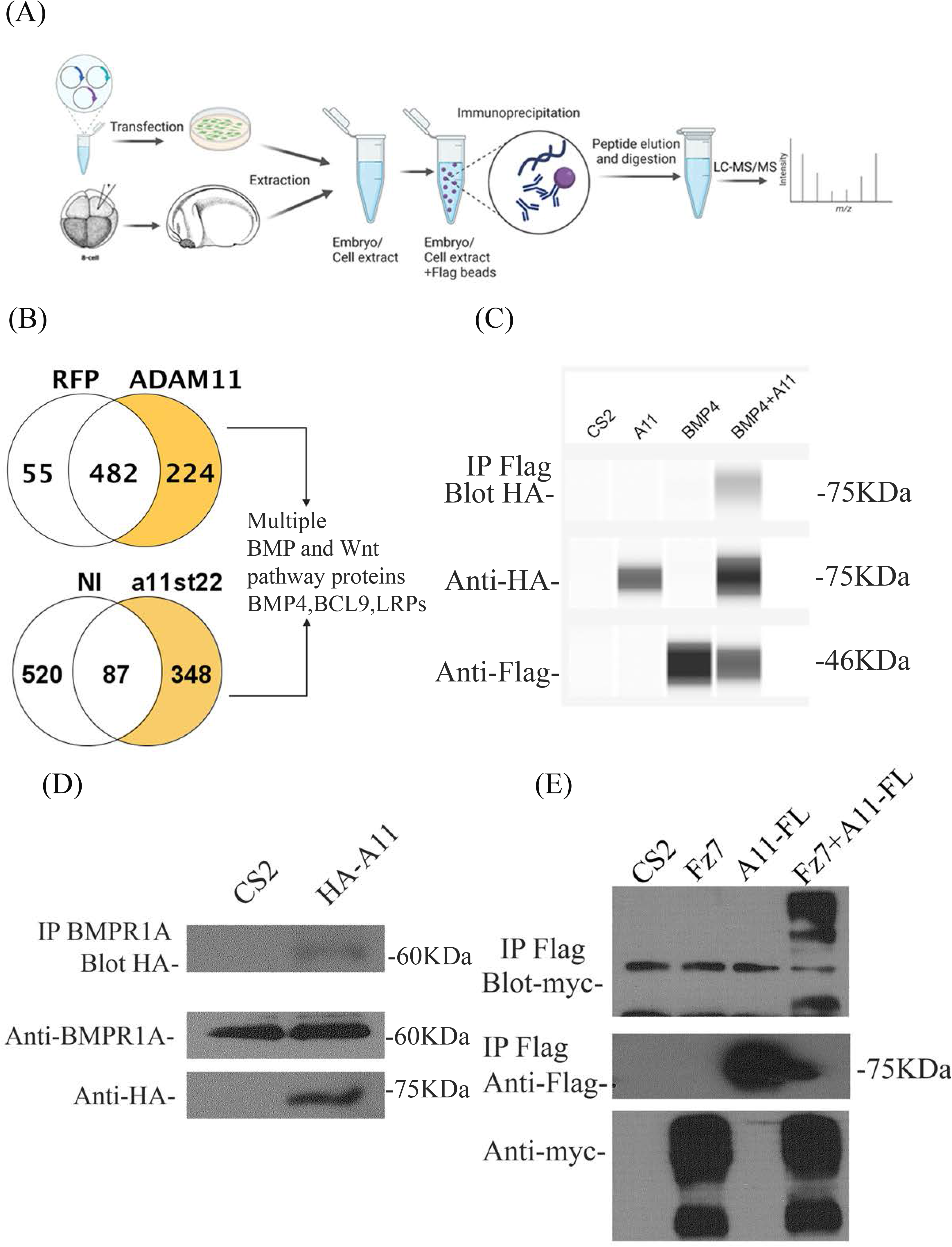
Adam11 binds to proteins of the BMP4 and Wnt signaling pathways. A) Schematic representation of the Lc/Ms/Ms experiment. Both human Hek293T cells and embryos were used to produce Adam11L-Flag. Hek293T cells were transfected while embryos were injected at the 8-cell stage in a dorsal animal blastomere to target CNC with Adam11L-flag mRNA or an irrelevant flag-tagged protein (RFP-Flag). The embryos were grown until stage 22 sorted and extracted. The Flag tagged proteins were immunoprecipitated and subjected to protein digestion and LC/MS/MS (Triplicate). (B) Proteins identified in either Xenopus embryos (348) or Hek293T cells (224) with at least two unique peptides that were absent in the negative controls were compared. (C) BMP4 binding to ADAM11 was confirmed by co-immunoprecipitation in Hek293T cells. HA-Adam11 was co-transfected with BMP4-Flag. Immunoprecipitation was performed using an anti-Flag antibody and western blot performed with both anti-HA and anti-Flag antibody. (D) Endogenous human BMPR1A binding to Xenopus ADAM11 was confirmed by co-IP in Hek293T cells. HA-ADAM11 or empty CS2 vector were transfected. Immunoprecipitation was performed using an anti-BMPR1A antibody and western blot performed with both anti-HA and anti-BMPR1A antibody. (E) Frizzled7 binding to ADAM11 was confirmed by co-IP in Hek293T cells. Adam11-Flag was co-transfected with Frizzled7-myc. Immunoprecipitation was performed using an anti-Flag antibody and western blot performed with both anti-Flag and anti-myc antibody.

### Loss of Adam11 increases Hsp90ab1 expression

To understand better how the Adam11 KD changed the state of the CNC cells we performed mass spectrometry on the dissected CNC explants (Fig.7A). After analysis of triplicate experiments, while the majority of proteins identified were common (585) we found 40 that were lost in the morphant and 80 that were present only in the CNC KD for Adam11. In particular, we found that Hsp90ab1 was only detected in Adam11 KD CNC. Hsp90ab1 has been shown to interact with the β-catenin destruction complex (Cooper et al., 2011) as well as LRP5 (Wang et al., 2019) protein to increase β-catenin signaling and promote EMT. To confirm the result from the Mass spectrometry experiments, we performed immunofluorescence on CNC explants using an antibody to Hsp90ab1 (Fig.7B-C). These results are strikingly similar to those observed for CyclinD1 (Fig.4B) and confirm the upregulation of Hsp90ab1 in CNC lacking Adam11.

**Figure 7:**
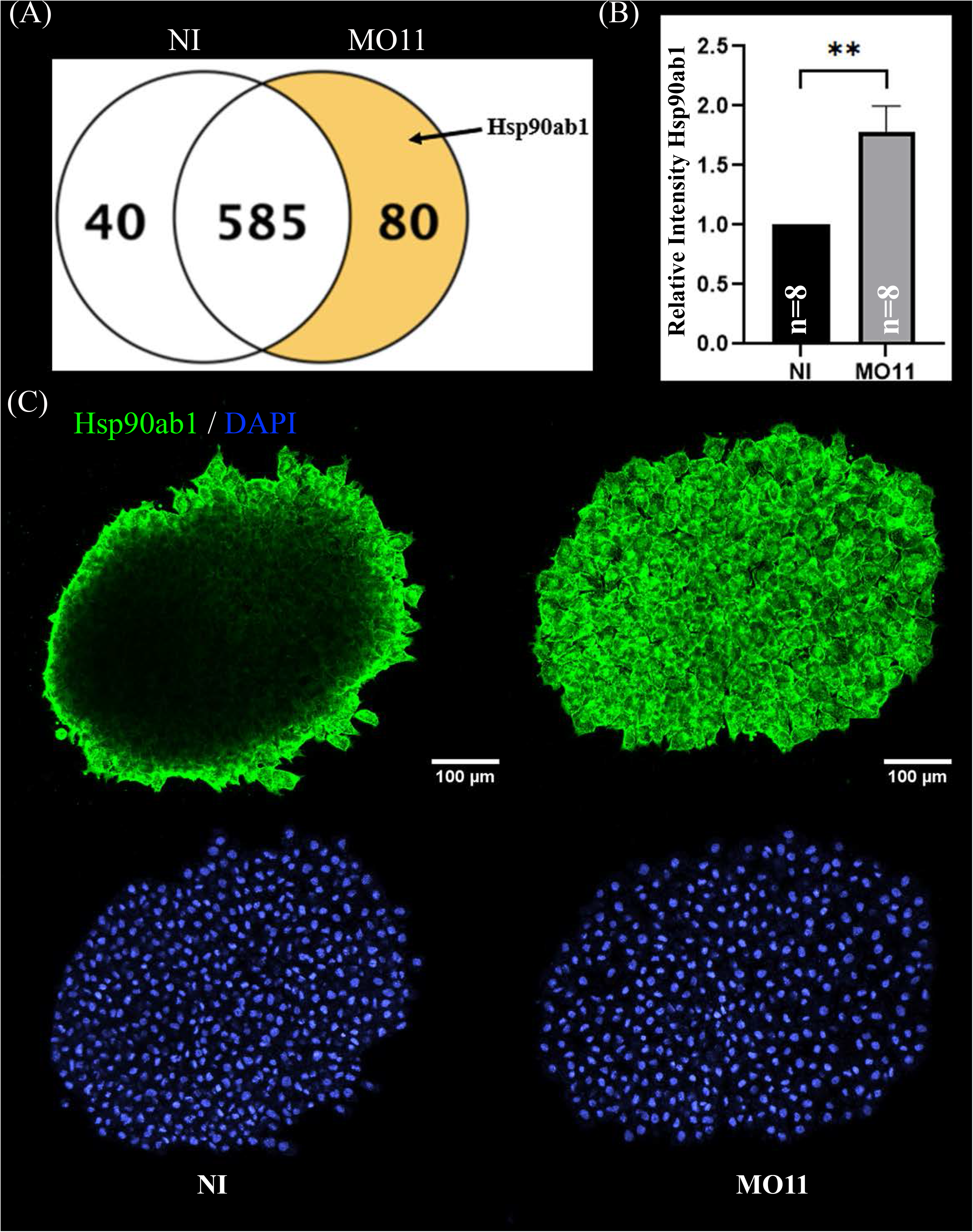
Adam11 KD increases Hsp90ab1 expression. (A) Ven diagram of proteins expressed in CNC explant from non-injected control embryos (NI) or embryos KD for Adam11 (MO11). B) Relative fluorescence intensity of Hsp90ab1 in CNC explants shown in (C). (C) Confocal imaging of CNC explants one hour after dissection. CNC explants were dissected at stage 17 and placed on fibronectin coated glass coverslip. Explants were stained for Hsp90ab1 (green) and DAPI (Blue). Students t-test was done for statistical analysis (**=P<0.01, three independent experiments).

### Adam11 increases BMP4 signaling

As shown above, Adam11 interacts with both BMP4 and its receptor BMPR1A (Fig.6). We also found that secreted BMP4 interacted with shed Adam11 in the conditioned media of co-transfected cells (data not shown). We therefore tested if Adam11 binding to BMP4 increased or decreased BMP4 signaling. For this, we used the animal cap (AC) assay. The intact animal cap represents a naive ectoderm with high BMP4 signaling and no expression of Adam11 based on published RNAseq data (Haas et al., 2019; Sive et al., 2007; Xu et al., 1995). Dissociation of the animal cap dilutes BMP4 signaling and induces neural tissue that expresses a high level of Sox3 (Fig.8A). We repeated these experiments using AC expressing injected Adam11L. Following animal cap dissociation into single cells, we found that Sox3 was significantly reduced in explants from embryos overexpressing Adam11 (Fig.8A, A11AC). In the same samples, we found that the phosphorylated Smad1/5 was increased by Adam11 overexpression (Fig.8B), consistent with an activating role of Adam11 toward BMP4. To further test the activity of Adam11 towards BMP4 signaling during neurulation, we performed a luciferase assay using the pGL2-15xGCCCG-lux BMP reporter (Alkobtawi et al., 2021) targeted at the 8-cell stage (Fig.8C). The luciferase activity was decreased with Adam11 KD further confirming that Adam11 increases BMP4 activity. BMP4 plays a critical role during neural induction by defining the ventral limit of the neural tissue, while BMP4 inhibitors are essential for neural induction (Weinstein and Hemmati-Brivanlou, 1997). To test if Adam11 interaction with BMP4 was important for the neural tube closure phenotype (Fig.1C), we repeated the assay using low doses of BMP4 mRNA (12.5 pg) to rescue the neural tube closure phenotype (Fig.8D-E). Indeed, injection of BMP4 in the dorsal part of the embryo rescued the neural tube closure timing to a level indistinguishable from the non-injected control (Fig.8E). Taken together these results show that Adam11 interacts with the BMP4 ligand and receptor to increase local signaling to a level critical for proper neural tube closure. The role of Adam11 in controlling BMP4 signaling levels is also consistent with our observation that Adam11 KD increases the domain of expression of neural markers such as *sox2* and Sox3 (Fig.3).

**Figure 8:**
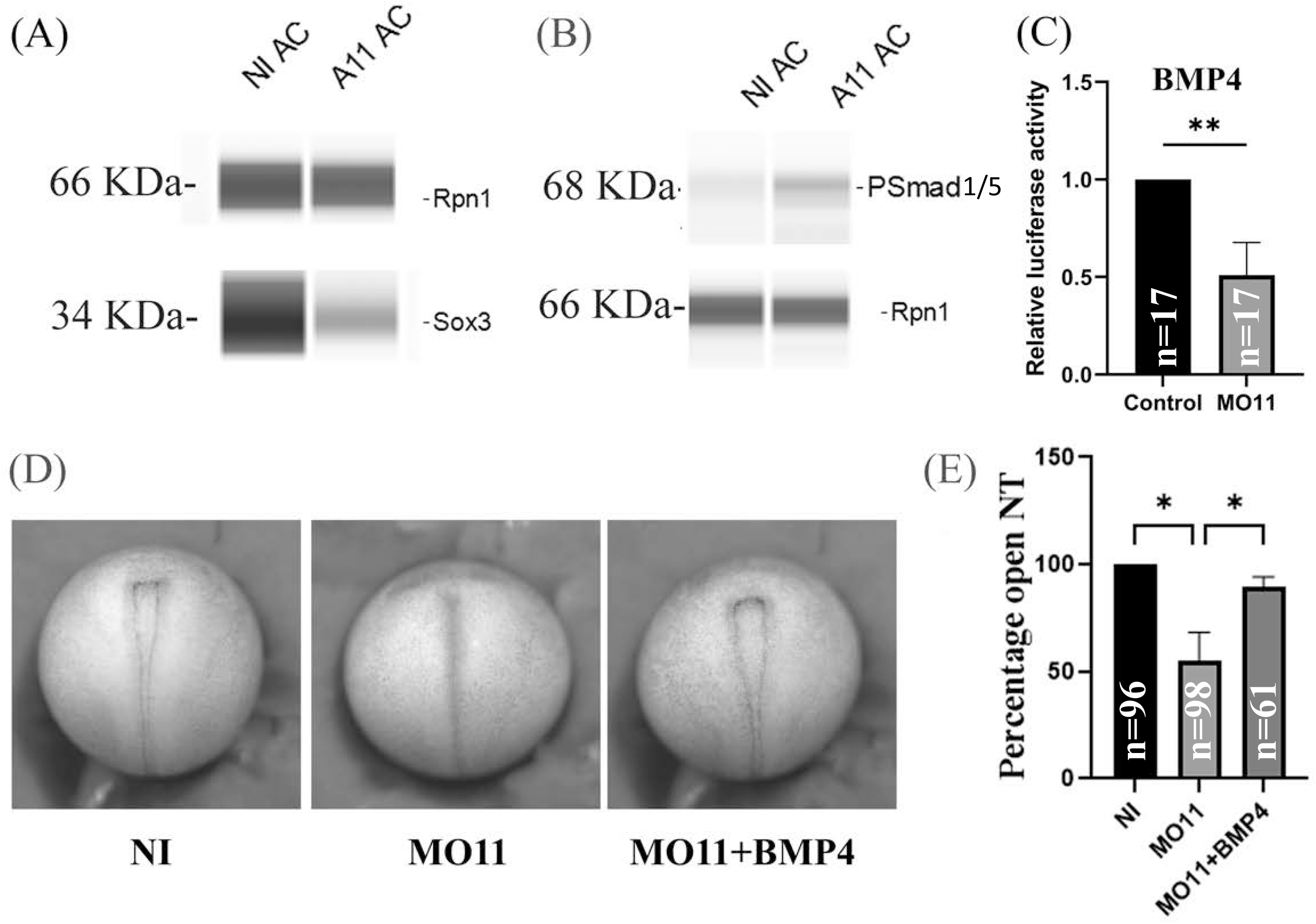
Adam11 increases BMP4 signaling. (A-B) Capillary western blot. Animal cap from control embryos or embryos injected with Adam11 mRNA (1 ng) were dissected at stage 9, dissociated in CMF for 2 hours and reassociated and grown in Danilchick media until sibling embryos reached stage 20. Proteins were then extracted and analyzed by capillary western blot using antibodies to ribophorin1 (rpn1) as a loading control and either Sox3, to detect neural induction (A), or phospho-smad1/5 antibody to measure BMP4 activity (B). (C) Histogram representing the relative luciferase activity between control and Adam11 KD. Embryos were injected at 8-cell stage with 1.5 ng of MO11 and 20 pg of pGL2-15xGCCCG-lux BMP reporter and 4 pg of CMV renilla (p<0.01). (D) Representative photograph of neurula stage embryos (Dorsal view anterior up). Embryos were injected at the one cell stage with either MO11 or MO11 and 12.5pg of BMP4 mRNA. (E) Histogram representing statistical analysis of the phenotype represented in D. Asterisk represent the statistical significance at p<0.05 (*) p< 0.01 (**).

### Adam11 expression in cancer

We have shown that Xenopus Adam11 restricts β-catenin activity in the CNC. Loss of Adam11 increases single CNC cell migration *in vitro* resembling an epithelium to mesenchyme transition (EMT). Given the role of β-catenin in cancer and the known relation between EMT and cancer metastasis, we looked at human *ADAM11* gene expression in available databases (Bartha and Gyorffy, 2021). Out of 56,938 unique multilevel quality-controlled samples, *ADAM11* expression was found to be significantly different in 18 out of 23 tumor types (Fig.9A, red). It was significantly reduced in 16 of the 18 tumor types. Given the previous observation that *ADAM11* was mutated in two breast cancer samples (Emi et al., 1993), we further focused our analysis on data obtained from invasive breast carcinoma (Fig.9B). The expression level was significantly lower in the samples from cancer patient (1097 patients) when compared to normal tissues (403 patients, mean FC=0.35, p=10^-84^). Interestingly in the same samples, *CYCLIND1* was found to be significantly increased (mean FC= 4.15, p=2.29 10^-76^). Taken together these results strongly suggest that Adam11 ability to restrict β-catenin activity and delay EMT in the CNC is conserved in adult tissues and that loss of ADAM11 may contribute to β-catenin hyper activation, and *CYCLIND1* overexpression, leading to tumor progression and metastasis.

**Figure 9:**
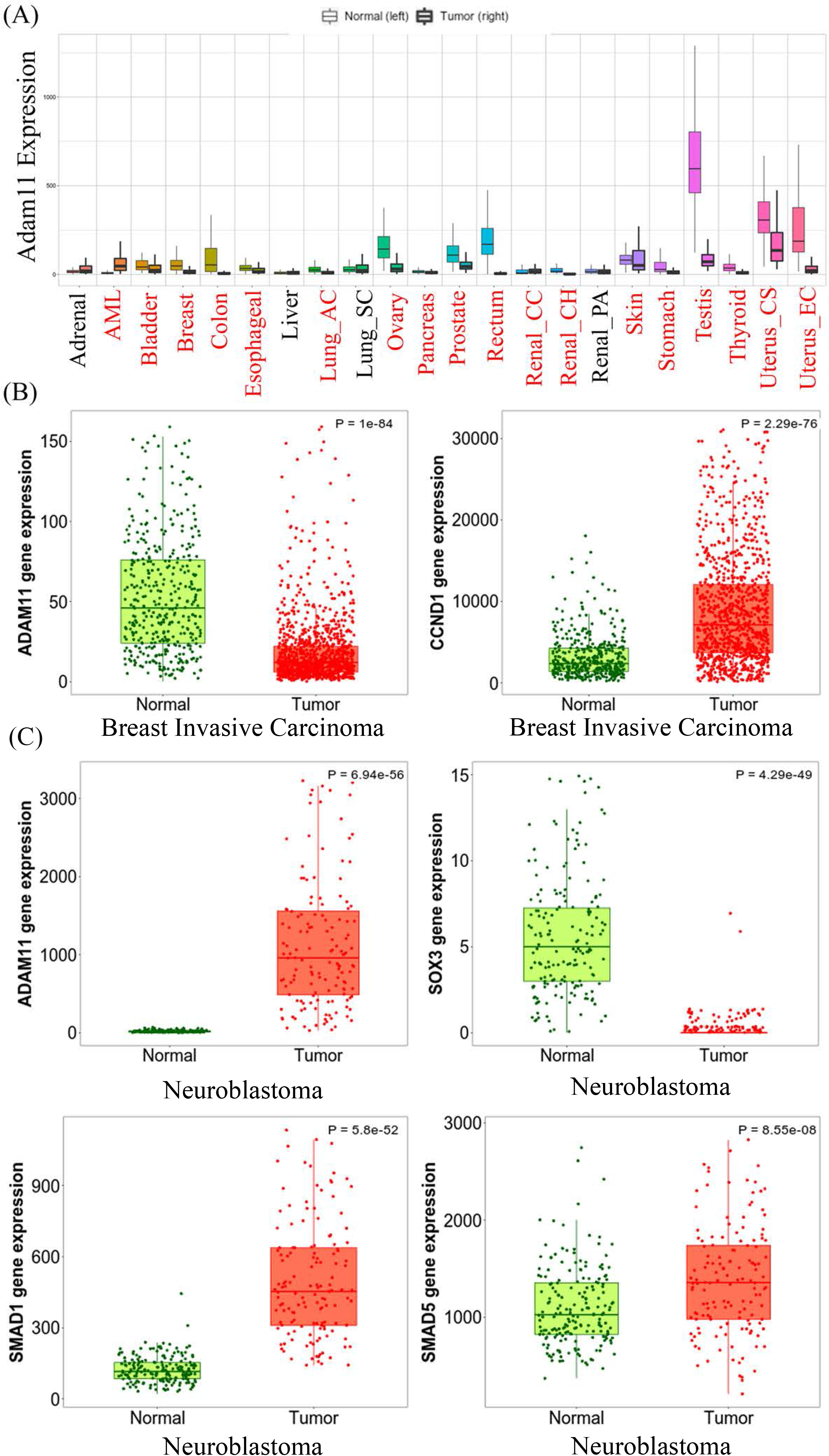
ADAM11 expression in cancer and cancer metastasis. Tnm plot analysis of *ADAM11* expression levels between normal tissue samples and tumors pan-cancer from human patients. Tumor types with significant changes in *ADAM11* expression are indicated in red. (B) Tnm plot analysis of *ADAM11* and *CYCLIND1* expression between multiple normal (403) and tumor (1097) breast invasive carcinoma patient samples. (p=10^-54^ and p=2.29 10^-76^ respectively). (C) Tnm plot analysis of *ADAM11*,(p=5.94 10^-56^), *SOX3*, (p=4.29 10^-49^), *SMAD1* (p=5.8 10^-52^), and *SMAD5* (p=8.55 10^-5^) expression between Neuroblastoma (149) and normal (190) patient samples. Mann-Whitney test was done for the p-values indicated in the legend.

One type of tumor in which *ADAM11* is increased rather than decreased is neuroblastoma (Fig.9C, 190 normal and 149 tumor samples, mean FC=61.52 p=6.94 10^-56^). In these tumors, we found that *SMAD1*and *SMAD5* were significantly increased (mean FC=4.46 p=5.8 10^-52^ and FC=1.31 p=8.55 10^-8^ respectively) while *SOX3* was significantly decreased (mean FC=0.13 p=4.29 10^-49^). An observation that is consistent with our results showing that the overexpression of Adam11 in the naïve animal cap ectoderm increases BMP4 signaling resulting in increased Smad1-5 phosphorylation and decreased Sox3 (Fig.8), while loss of Adam11 increases Sox3 expression (Fig.3E-F).

### Reduction of ADAM11 in B16 melanoma increases cell proliferation

The human cancer analysis points to a strong correlation between *ADAM11* expression and *CYCLIND1* levels. To further test if *Adam11* expression in cancer cells could modulate the Wnt signaling pathway, we used mouse B16 melanoma cells (Poste et al., 1980). These cells are derived from neural crest cells, express a relatively high level of *Adam11*, and are highly invasive *in vivo* when injected in the blood stream but not subcutaneously. Furthermore, in skin cutaneous melanoma, published RNA-seq data shows a decreased level of *ADAM11* and an increased level of *CYCLIND1* between tumors (103 patients) and normal samples (474 patients, Fig.10A). We used shRNA mediated inhibition of Adam11 as well as rescue with Xenopus Adam11 in these cells to test the effect on cell proliferation. Based on the CNC results, we expected that loss of Adam11 would increase Wnt signaling and CyclinD1 expression (Fig.4) resulting in an increased cell proliferation (Fig.5). While we obtained only a modest but reproducible reduction in *Adam11* expression (35%, Fig.10B), we were able to see an increase in the level of β-catenin by western blot (Fig. 10C). We also found that this reduction of *Adam11* in B16 increased the cell proliferation rate by nearly 40% (Fig.10D). We then tested by RT-qPCR the expression of multiple markers of EMT and target of β-catenin. We observed a statistically significant increase in *CyclinD1* and *Snail* levels (Fig. 10E), and while there was a small decrease in *E-cadherin* and increases in *N-cadherin* and *Vimentin* that fit with our EMT expectation those were not significant. In order to understand the increase in cell number (Fig.10D), we used the well characterized FUCCI cell cycle reporter (Sakaue-Sawano and Miyawaki, 2014), transfected in the context of Adam11KD and rescue using the Xenopus Adam11L construct and used FACS to follow cell cycle progression. Following successful transfection, the cells are red when they are in G1, yellow during the transition from G1 to S or green if they are in S, G2 or M phase (Fig. 10F). We found that the decrease in *Adam11* resulted in a significant decrease in the number of cells in the G1-S transition and increased number of cells in S, G2 and M phases (Fig.10G). The differences obtained with the silencing of *Adam11* was partially rescued using Xenopus Adam11, but the variability was to high for significance (Fig.10G).

**Figure 10:**
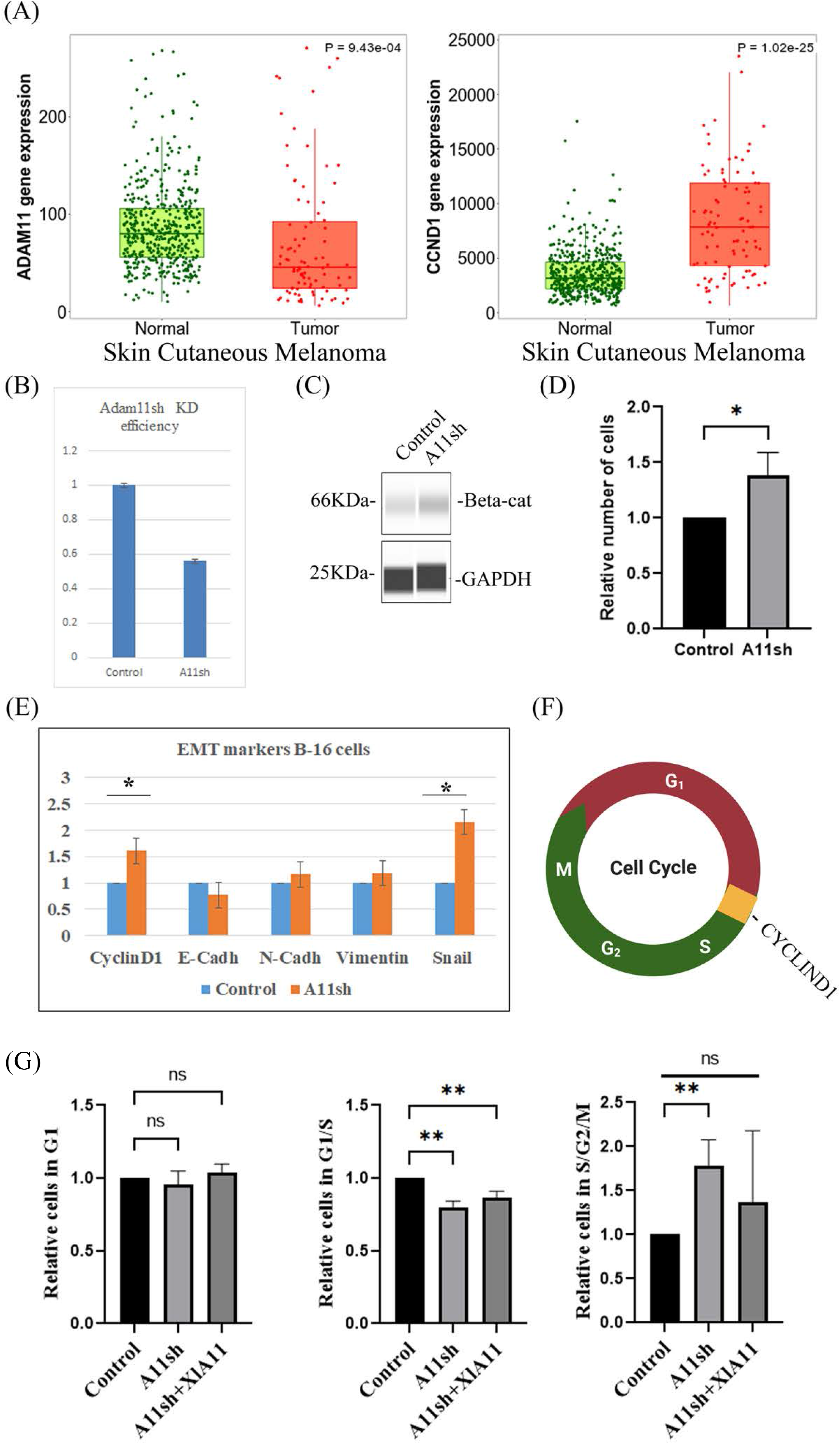
ADAM11 expression in Melanoma. Tnm plot analysis of *ADAM11* (p=9.43 10^-4^) and *CYCLIND1* (p=51.02 10^-25^), expression between multiple normal (474) and tumor (103) skin cutaneous melanoma patient samples. (B) RT-qPCR analysis of mouse B-16 melanoma cells transfected with either empty plasmid control or A11shmouse Adam11sh plasmid, cells were harvested for mRNA and processed for RT-qPCR. *Adam11* levels between the samples was normalized to *Beta-actin.* (C) Capillary Western Blot of B-16 melanoma cells transfected with either empty plasmid control or Adam11sh (A11sh) plasmid, stained for β-catenin and Gapdh. (D) Histogram showing the relative number of cells present 48 hours post transfection in either empty plasmid control or A11sh plasmid condition. (E) RT-qPCR analysis of B-16 melanoma cells transfected with either empty plasmid control or A11sh plasmid. *Cyclind1, E-cadherin, N-cadherin, Vimentin, and Snail* levels between the samples was normalized to *Beta-Actin.* The fold change calculated using the ΔΔCT is indicated. Error bars represent the standard deviation (* means p<0.05). (F) Diagram showing the phases of the cell-cycle and the fluorescence they would emit in each phase after transfection with the FUCCI reporter plasmid. (G) Histogram depicting the relative number of cells in each phase following KD of mouse Adam11 (A11sh) and rescue with Xenopus Adam11 (XlA11) protein. The cells transfected with the empty vector were set to one for each phase. Cell numbers were obtained by FACS analysis (50,000 cells counted per experiment, 3 independent experiment, ** means p<0.01). As expected, given the regulation of CyclinD1, significantly less cells were found in the G1 to S transition and more cell in S, G2 and M.

## Discussion

Adam11 is a member of the non-proteolytic ADAM that appear essential for the proper function of the central nervous system in mammals (Takahashi et al., 2006a; Takahashi et al., 2006b). In addition, ADAM11 was identified as a putative tumor suppressor thirty years ago but no additional data in support of this hypothesis was ever published (Emi et al., 1993). Here we show that Adam11 controls Wnt and BMP4 signaling pathways during neurulation and neural crest cell migration. Loss of Adam11 increases β-catenin transcriptional activity and decreases BMP4 signaling. Loss of Adam11 accelerates neural tube closure and accelerates the transition from collective to single cell migration in the CNC. We further show that Adam11 expression is reduced in multiple tumors. We propose that Adam11 functions as a regulator of Wnt and BMP4 signaling to control the expression of neural and neural crest differentiation markers as well as the onset of EMT and morphogenetic movement in the embryo.

### ADAM11 in cancer

ADAM11 is well conserved between Xenopus Mouse and Human both in the extracellular and cytoplasmic domains. This conservation is especially striking in the short cytoplasmic domain that does not contain any putative functional attribute. The least conserved region is the pro-domain which in proteolytic ADAM controls the activation. Thirty years ago, ADAM11 was found to be mutated in two mammary gland tumors and was postulated to be a tumor suppressor (Emi et al., 1993). Surprisingly no other study tested this hypothesis. Given our results showing that Adam11 regulates β-catenin activity in Xenopus embryos (Fig.4), we analyzed the expression of *ADAM11* in available tumor expression data. We found that consistent with the tumor suppressor hypothesis *ADAM11* expression is reduced in the majority of solid tumor (Fig.9A). When focusing only on breast invasive carcinoma which were originally associated with *ADAM11/MDC* mutation and Melanoma which are neural crest derived. We find that in addition to the significant decrease in *ADAM11* (Fig.9 and 10), there was also a significant increase in *CYCLIND1*. While there is no causal relation in this observation, our data from Xenopus embryos and mouse B-16 melanoma cells shows that decreasing Adam11 protein level is sufficient to increase β-catenin activity and CyclinD1 expression suggesting a conserved function of Adam11 in cancer. It will be of interest to test if human ADAM11 can also regulate β-catenin and rescue neural tube closure timing and CNC migration in Xenopus embryos. On the other hand, it would be interesting to test if increasing ADAM11 protein level in cancer cells is sufficient to reduce β-catenin activity, *CYCLIND1* expression and cell proliferation. While this seems like an interesting strategy, we have also found that ADAM11 expression is increased in Neuroblastoma, a childhood cancer thought to be derived from neural crest cells. In neuroblastoma, the increase in ADAM11 expression is associated with increased expression of Smad1 and 5 and decreased expression of Sox3. This is similar to what we observed when overexpressing ADAM11 in naïve ectoderm in Xenopus embryos (Fig.8) suggesting that in Neuroblastoma ADAM11 positive regulation of BMP4 signaling could lead to the repression of neural differentiation markers (Sox2 and 3) resulting in more naïve cells. These naïve or “stem” cells could contribute significantly to the malignancy. Therefore, our observation suggests that a fine control of ADAM11 expression is critical for normal development as well as tissue homeostasis. A careful structure/function analysis will be required to understand the role of each of the functional domains to the overall function of ADAM11.

### Neural crest cell migration

We have found that CNC migrated well *in vitro* but not *in vivo*. This is certainly not unique as we found this to be the case in most ADAM inhibition highlighting the essential role of ADAM interaction with the extracellular environment (Cousin et al., 2012; McCusker et al., 2009). While β-catenin is essential for proper CNC induction, studies have shown that β−catenin activity needs to be reduced for proper migration and that abnormal activation within the CNC leads to inhibition of migration (Maj et al., 2016). This suggest that the excess activation of β−catenin following Adam11 KD could be responsible for the perturbation of CNC migration *in vivo*. In contrast both chemical inhibition of GSK3 and activation of TcF3 also inhibit migration *in vitro* preventing single cell dispersion (Maj et al., 2016). This is the opposite phenotype that what is observed in the Adam11 morphant embryos where single cells increase and both speed and persistence are higher. This suggest that the loss of Adam11 is not simply a hyper activation of β−catenin and that the decrease in BMP4 signaling may also influence CNC migration. Certainly, the activation of β−catenin following the loss of Adam11 is much more subtle as the overall level of the protein is unchanged and only the nuclear localization is increased.

Adam11 knock out mice show normal mechanical and noxious heat responses but have significantly decreased chemical nociception indicating that Adam11 is necessary for the formation or the relay of specific set of sensory information (Takahashi et al., 2006a). The sensory neurons are derived mostly from the cranial placodes and cranial neural crest (cranial ganglia) anteriorly and from the trunk neural crest posteriorly (Vermeiren et al., 2020). The loss of specific type of nociception in Adam11 knock out mice (Takahashi et al., 2006a) raises the possibility that the defect in these neurons might be a consequence of abnormal specification or migration of the neural crest. Adam11 transcripts have been detected in the dorsal root ganglia and the neural crest in the mouse (Diez-Roux et al., 2011; Rybnikova et al., 2002) and the frog (Cai et al., 1998). While no obvious anomalies were reported in the DRG of Adam11 KO mice, it is likely that the absence of the small nociceptive neurons would not be detected without using cell specific markers. Alternatively, it is also possible that these neurons exist but fail to properly organize their synapse as shown for the cerebellar basket cell (Kole et al., 2015). Further analysis of the Adam11 morphant embryos at later stage could help identify subtle DRG defects.

### Neural tube closure

Neural tube closure is a complex morphogenetic event that involves hundreds of different genes. In Xenopus, like in mammals, the neural folds elevate toward the center of the neural plate and then fuse. Based on asymmetric injection we find that the neural tube elevates and comes to the midline faster on the side missing Adam11 (data not shown). It is important to note that the neural plate, as defined by Sox3 expression, is wider on the side missing Adam11. Defects in the non-canonical Wnt planar cell polarity (PCP) pathway induce wider neural plate but contrary to Adam11 KD the neural fold fail to reach the midline (Goto and Keller, 2002; Wallingford and Harland, 2002). While the role of the PCP pathway is clearly critical, β-catenin has been shown to regulate the expression of Pax3 and Cdx2, two transcription factors critical for neural tube closure. In the absence of β-catenin, neural tube closure is inhibited but can be rescued by the expression of Pax3 (Zhao et al., 2014). Therefore, increase or early activation of β-catenin in Adam11 morphant embryos could lead to the activation of Pax3 and the early neural tube closure. In mouse, FGF3 mutants have elevated levels of BMP signal, increased neural cell proliferation and delay neural tube closure (Anderson et al., 2016). This would be consistent with the decreased BMP4 signaling and accelerated neural tube closure in Adam11 morphant embryo and could explain the rescue observed using low levels of BMP4 mRNA (Fig.8D). Careful analysis of β-catenin and BMP4 signaling levels prior to neural tube closure would be critical to determine if they are indeed responsible for this Adam11 phenotype. In the future, it would be interesting to determine if the early closure of the neural tube affects the proper specification of the various neuronal cell type and their organization. This could in part explain the neurological disorder observed in mice lacking Adam11. It is clearly much more difficult to evaluate neural tube closure timing in mouse embryos and as such it is understandable that those have not been reported.

### Model (Fig.11)

We have shown that Adam11 binds to BMP4 and its receptor BMPR1A (Fig.6). Our results using the animal cap dissociation experiment (Fig.8) show that Adam11 increases BMP4 signaling, possibly by retaining BMP4 at the plasma membrane and presenting the protein to the receptor (Fig.11). Thus, Adam11 expression at the surface of the CNC would allow them to amplify the gradient of BMP4 locally restricting the expression of neural markers such as Sox2 and Sox3 to define the CNC zone (Fig.3). At the same time, Adam11 can also bind to Wnt receptors Fzd as well as multiple LRP proteins (Fig.6). We propose that the interaction of Adam11 with these Wnt receptor and co-receptor reduce their capacity to either bind to the Wnt ligand or to induce the cytoplasmic cascade that inhibit the destruction complex of β-catenin (Fig.11). Given the capacity of several ADAM protein to bind to the cysteine rich domain of Fzd or Sfrp proteins (Abbruzzese et al., 2015; Esteve et al., 2011), the usual binding site of the Wnt ligand (Dijksterhuis et al., 2015), Adam11 could simply act as a competitive inhibitor of the Wnt/Fzd complex. Alternatively, Adam11 could wedge itself between the Fzd and LRP proteins reducing the activation of the complex. In CNC lacking Adam11 we find a much higher Hsp90ab1 protein expression. This protein has been shown to stabilize Lrp proteins thus reinforcing the Wnt signaling pathway. Refining the composition of Adam11 protein complex and the binding sites of each protein will be required to understand the exact mechanism of Adam11 inhibition of β-catenin activation.

**Figure 11:**
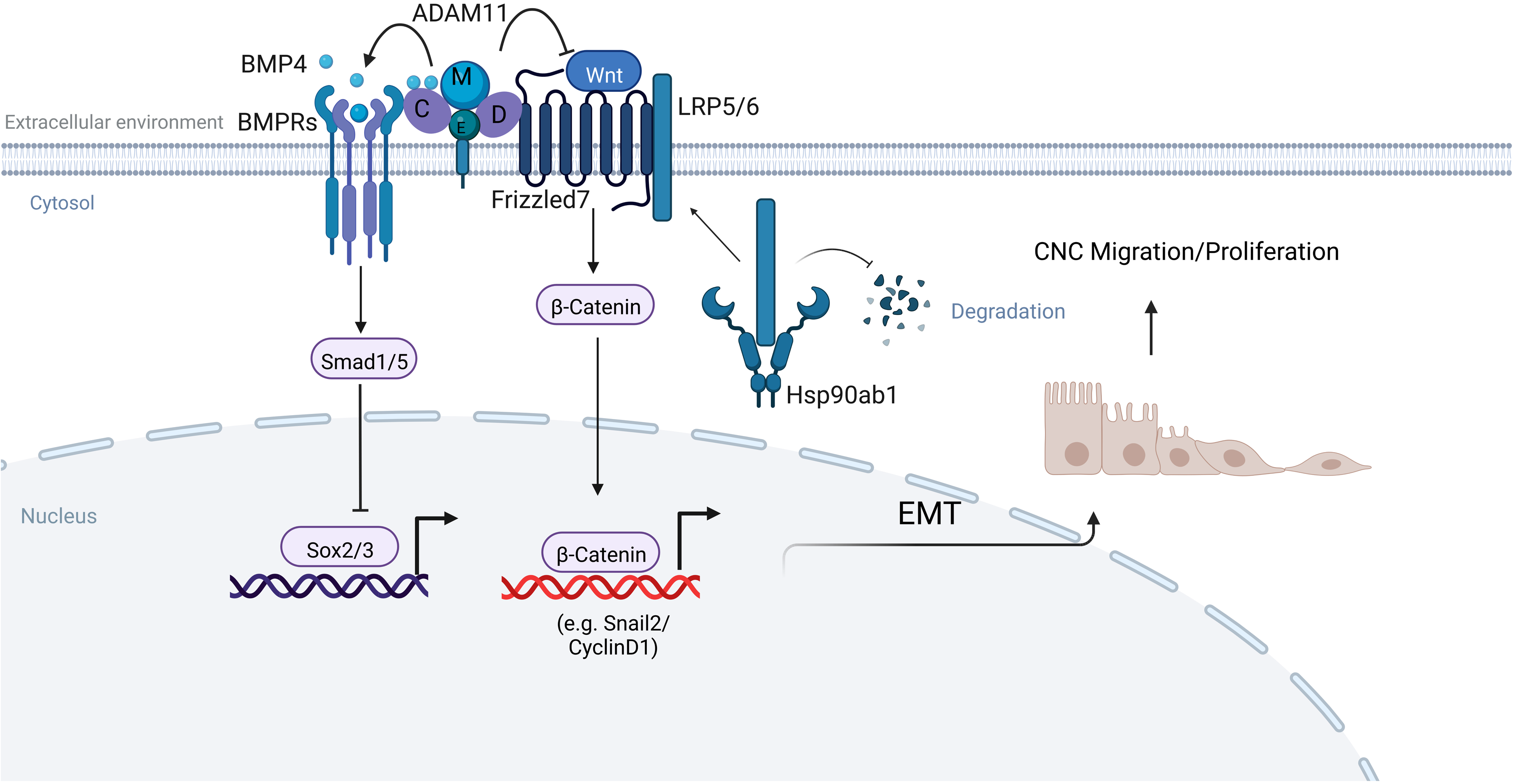
Model of Adam11 function. Adam11 binds to BMP4 and Frizzled on the plasma membrane to increase BMP4 and decrease Wnt/β-catenin signaling. The result of the increase BMP4 signaling is the inhibition of the neural marker Sox2 and Sox3. The result of the decreased in β-catenin activity is a reduction is CyclinD1 and Slug/Snail2 leading to less cell proliferation and delayed EMT. Loss of Adam11 in CNC leads to an increase in Hsp90ab1 that is known to stabilize Lrp5/6 proteins thus increasing the Wnt receptor signal and β-catenin translocation to the nucleus. Given the interaction of Adam11 with Fzd and Lrp, it is likely that the increase in Hsp90ab1 is a feedback loop reinforcing the pathway rather than the initial trigger of β-catenin activation. Adapted from “Signaling Pathways Underlying EMT and EndMT: Emerging Roles in Age-related Macular Degeneration (AMD)”, by BioRender.com (2022). Retrieved from https://app.biorender.com/biorender-templates

## Acknowledgment

This work was supported by grant from the NIH to DA (R24-OD21485), and DA and HC (R01DE016289). We thank members of the Alfandari lab for helpful comments, discussion and corrections. Relative contribution: Conceptual design (AP and DA). AP performed most of the experiments, BH produced the Xenopus antibodies used in this study and contributed to the cell culture work. DA perform CNC dissections and the proteomics experiment and analysis; HC performed the grafts and part of the targeted injections. AP and DA wrote the manuscript.

## Materials and Methods

### Antibodies

The following antibodies were used in this study: monoclonal antibody to Ribophorin-1 (Rpn1) was used as loading control, RRID:AB_2687673 (Khedgikar et al., 2017). The rat monoclonal to flag tag (Agilent Cat# 200474, RRID:AB_10597743). The monoclonal to *Xenopus laevis* Sox3 (DSHB Cat# DA5H6, RRID:AB_2876376) was developed in the laboratory (Horr et al., 2023) against the full length *Xenopus laevis* protein produced in Hek293T cells. It has been characterized by Western blot and immunofluorescence (R24OD21485). The rabbit polyclonal to HA tag (Applied Biological Materials Cat# G166, RRID:AB_2813867). The Rabbit monoclonal to phospho-Smad 1/5 (Cell Signaling Technology Cat# 9516, RRID:AB_491015). Mouse monoclonal to BMPR1A (Thermo Fisher Scientific Cat# MA5-17036, RRID:AB_2538508). Rabbit polyclonal to β-catenin (Abcam Cat# ab16051, RRID:AB_443301). Rabbit polyclonal to Cyclin-D1 (Bioss Cat# bs-0623R, RRID:AB_10856925). Mouse monoclonal to Hsp90ab1 (Thermo Fisher Scientific Cat# 37-9400, RRID:AB_2533349). The anti-myc antibody 9E10 (DSHB Cat# 9E 10, RRID:AB_2266850). Rat monoclonal against N-cadherin (DSHB Cat# MNCD2, RRID:AB_528119). Mouse monoclonal to *Xenopus laevis* Adam11.L (3C5) was developed in the lab by injecting a bacterial fusion protein containing the DC (disintegrin and cystein rich) domain and isolating the clone by the methods described in (Horr et al., 2023).

### Morpholinos and DNA constructs

Morpholino antisense oligonucleotides (Gene Tools, Philomath OR) were designed to either block the translation of Adam11L and S (MO11 Fig.1, CATCATTCATACATAATCATTCAGC) or the splicing of Adam11L (MO11spl, CCAATGGAGGCATAGTGGCACTG). Adam11.L was cloned in pCS2+ using cDNA made from RNA extracted from stage 18 X*enopus laevis* embryos. Flag and HA tags were added using Takara infusion cloning according to the manufacturer’s instructions. BMP4-Flag in pCS2 was a generous gift from Dr. Chenbei Chang (University of Alabama). Top Flash was previously described in (Chuang et al., 2010). All constructs were sequenced and transfected in 293T cells to validate protein translation, size and proper cell compartment expression.

### Injections and microdissections

SP6 polymerase was used for capped mRNAs synthesis after digesting using Not1 (Cousin et al., 2011). Injectors were calibrated using a 1 µl capillary needle (Microcaps, Drumond, PA, USA). The injection pressure was set at 15 psi and the injection time set between 50 and 200 ms to obtain a 5 nl delivery. Embryos were injected at 1-cell, 2-cell, 8-cell and 16-cells as described previously (Khedgikar et al., 2017). Embryos were raised at 14 to 15℃ until scoring for neural tube closure or CNC migration. For each injection, percentage of open NT was normalized to control embryos and was set to 100%, same was done for CNC migration using Mbc injected embryos. CNC grafts were performed as described (Alfandari et al., 2001; Cousin, 2018). CNC explants migration assays were performed as described in (Alfandari et al., 2003). All experiments were performed at least three times using different females to determine statistical significance.

### Cell culture and transfection

Hek293T (ATCC, CRL-3216 RCB2202) cells were cultured in RPMI media supplemented with 10 U/ml Pen/Strep, 2 mM L-glutamine, 0.11 mg/ml sodium pyruvate, and 10%FBS (Hyclone, South Logan, UT). Transfections were performed using Polyethylenimine (PEI, Polysciences Inc.). For each transfection, 1µg of DNA was mixed with 10µg of PEI in 200µl of Optimem media (Hyclone) at room temperature for 15min prior to adding to the cells dropwise. The media was changed after 24h post transfection.

Mouse melanoma B16/F10 (CRL-6475) cells were a generous gift from Dr. Leonid pobezinsky, the cells were cultured in DMEM media with 10 U/ml Pen/Strep and 10%FBS. Transfections were performed as described for Hek293T.

### Animal cap assays

Animal caps dissociation and reaggregation was done as described in (Sive, Grainger et al. 2007). Briefly, 10 to 20 animal caps were dissected in 1XMBS (1XMBS: 88.0 mM NaCl, 1.0 mM KCl, 2.4 mM NaHCO_3_, 15.0 mM HEPES [pH 7.6], 0.3 mM CaNO_3_-4H_2_O, 0.41 mM CaCl_2_-6H_2_O, 0.82 mM MgSO_4_) containing 50µg/ml of gentamycin. They were then transferred to Calcium, Magnesium free MBS (CMF) supplemented with 1mg/ml of BSA for 2h with gentle rocking to dissociate the animal cap into single cells suspension. The cells were then transferred into conical 96 well (Thermofisher) plate precoated with PBS 1% BSA and incubated overnight in Danilchik media (53 mM NaCl, 11.7 mM Na_2_CO_3_, 4.25 mM potassium gluconate, 2 mM MgSO_4_, 1 mM CaCl_2_, 17.5 mM Bicine, 1 mg/ml BSA, pH 8.3) until control embryos reached stage 20.

### Quantitative PCR

Quantitative real time PCR was done as previously described (Neuner et al., 2009). All primers were tested for efficiency. Embryos were collected at stage 15-18, B-16 cells were collected after 48 hours of transfection and RNA was extracted using (Roche, RNA isolation Kit). Total RNA was quantified on a nanodrop (Thermofisher) at 260nm. The cDNA was made using (Invitrogen Superscript III) according to manufacturer’s instructions. Quantitative PCR was performed using SYBR green (Takara, Kyoto, Japan) to measure mRNA levels of Adam11.L and GAPDH was used to normalize total cDNA amount, in case of B-16 cells beta-actin was used to normalize the total cDNA amount. The relative gene expression was calculated using the ΔΔCT method as described (Livak and Schmittgen, 2001). The result is represented in fold change compared to non-injected control. Oligos used are described in the table below (xl indicate Xenopus laevis primer, all other primer are mouse).

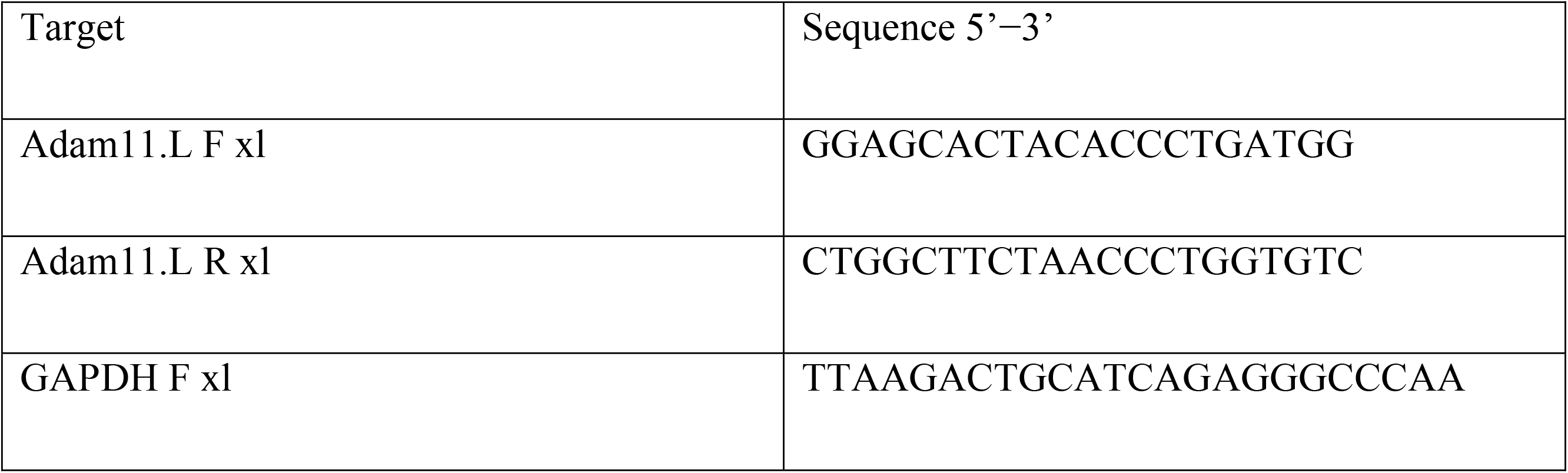
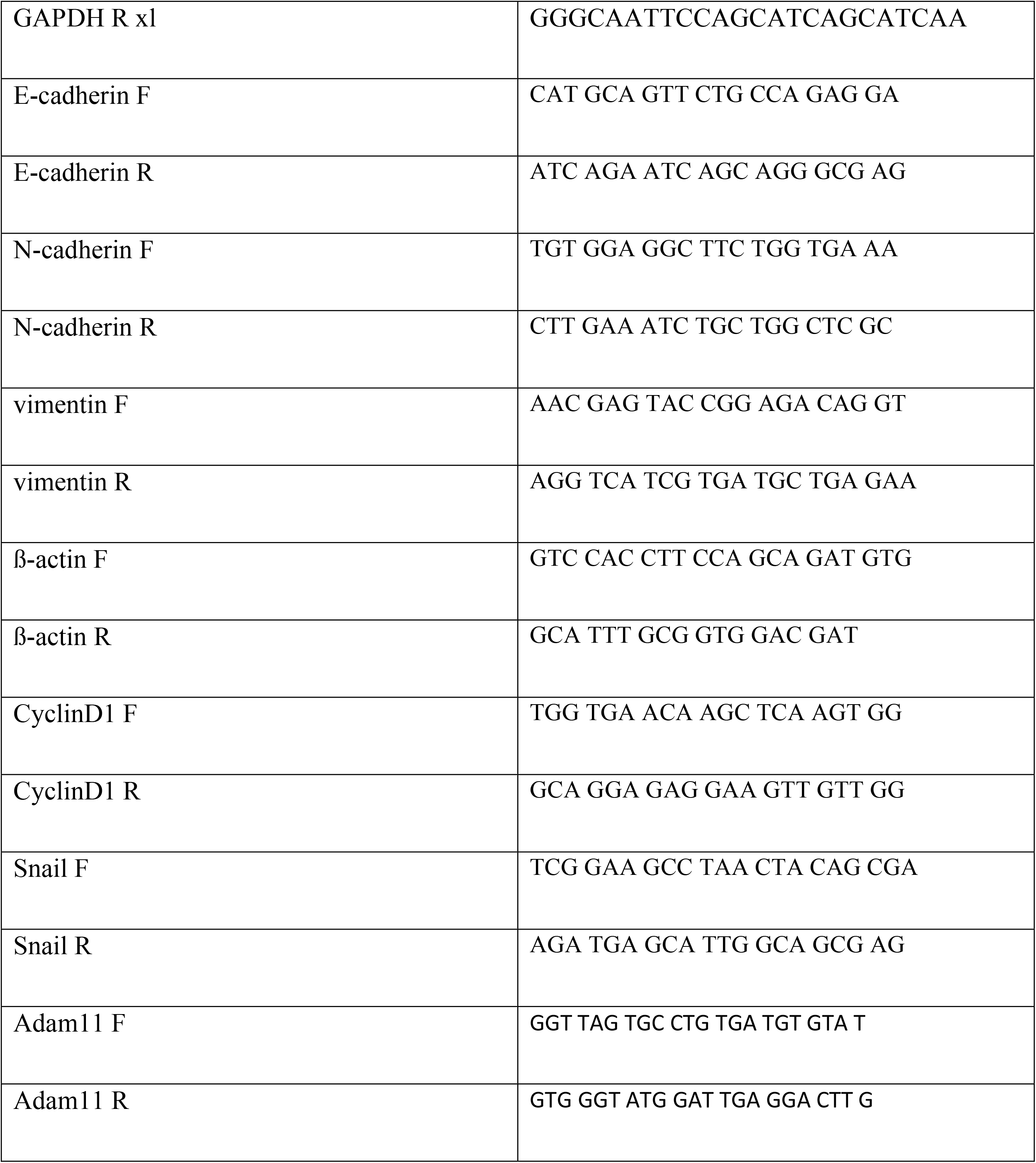

### Immunoprecipitation and western blots

Hek293T, embryos, Animal caps and CNC explants were extracted in 1XMBS-1% Triton-X100, Protease phosphatase inhibitor cocktail 1X (Thermoscientific) and 5mM EDTA. Immunoprecipitations were performed using 1-2ug of antibody bound to protein A/G magnetic beads (Thermofisher), incubated overnight at 4℃. The beads were washed 3 times for 5 minutes with extraction buffer at room temperature then eluted in 2X reducing laemmli sample buffer. All proteins were separated in 5-22% gradient SDS-PAGE gels and transferred to polyvinylidene fluoride membranes (PVDF, Millipore, Billerica, MA) using a semi-dry transfer apparatus (Hoeffer). Quantitative capillary western blots were performed using Proteinsimple automatic western machine (Wes) according to manufacturer’s instructions.

### Whole mount *in situ* hybridization

Whole mount *in situ* hybridization was done using previously described protocol (Harland 1991). The probes for *Sox2, Snai2/Slug, Sox9, Sox8* were synthesized using diogoxigenin-rUTP–label.

### Immunofluorescence

CNC cells were dissected at stage 17 and placed on fibronectin coated glass bottom plates (20ug/ml) for 1 to 3 hours at 18℃ (Cousin and Alfandari, 2018). The explants were fixed using MEMFA (0.1 M MOPS pH 7.4, 2 mM EGTA pH 8, 1 mM MgSO4 and 4% paraformaldehyde) for 1 hour, permeabilized using 0.5% TX100 in 1XMBS with 100mM glycine for 1 hour. The explants were then blocked using PBS containing 10% heat inactivated goat serum, 1% BSA, 0.1% Tween for 1h prior to incubation in the same buffer with the primary antibodies overnight at 4℃. CNC explants were washed in PTw (PBS-tween 0.1%) 3 times 15 min, blocked again for 1 hour at room temperature using blocking solution prior to incubation with the secondary antibody and Hoest33342 for 1h at room temperature. The explants were washed 3 times in PTw and imaged using Nikon confocal microscope (A1RHD25).

### Mass spectrometry

Protein extracted from 10 CNC explants as described above or immunoprecipitated with the Flag antibody from embryos or Hek293T cells were diluted in 8M urea and processed with Trypsin/Lysine-C according to manufacturer instruction (Promega). Tandem mass spectrometry was performed using a Thermo orbitrap Fusion (Mass spectral data were obtained at the University of Massachusetts Mass Spectrometry Core Facility, RRID:SCR_019063). All MS/MS samples were analyzed using Sequest (Thermo Fisher Scientific, San Jose, CA, USA; version IseNode in Proteome Discoverer 2.4.1.15). Sequest was set up to search XenbaseProteinXL2020.fasta (unknown version, 72266 entries) or the human protein database uniprot-human containing the sequences for Xenopus Adam11 (126358 entries) assuming the digestion enzyme trypsin. Sequest was searched with a fragment ion mass tolerance of 0.60 Da and a parent ion tolerance of 10.0 PPM. Carbamidomethyl of cysteine was specified in Sequest as a fixed modification. Oxidation of methionine, acetyl of the n-terminus and phospho of serine threonine and tyrosine were specified in Sequest as variable modifications.

Criteria for protein identification: Scaffold (version Scaffold_5.0.1, Proteome Software Inc., Portland, OR) was used to validate MS/MS based peptide and protein identifications. Peptide identifications were accepted if they could be established at greater than 95.0% probability by the Peptide Prophet algorithm (Keller et al., 2002) with Scaffold delta-mass correction. Protein identifications were accepted if they could be established at greater than 99.0% probability and contained at least 2 unique identified peptides. Protein probabilities were assigned by the Protein Prophet algorithm (Nesvizhskii et al., 2003). Proteins that contained similar peptides and could not be differentiated based on MS/MS analysis alone were grouped to satisfy the principles of parsimony. Proteins sharing significant peptide evidence were grouped into clusters.

### Flow Cytometry

Flow cytometry was performed on B-16F10 cells that were transiently transfected with the various plasmids. Only the cells successfully transfected with FUCCI were fluorescent and analyzed. The cells were harvested 36 to 48 hours post transfection using a cell scraper and resuspended in PBS 1% BSA. Flow cytometry was performed on a BD DUAL LSRFortessa and the data was analyzed using FACSDiva 8.0. 50,000 cells were acquired for each sample in three independent experiments.

